# Lineage-aware stochastic modeling reveals gene-expression dynamics in development and disease

**DOI:** 10.64898/2026.06.25.734628

**Authors:** Jiawei Xing, Stephen J. Staklinski, Zhihan Liu, Dawid Nowak, Adam Siepel

## Abstract

Gene expression changes along cell lineages, but most single-cell RNA-seq analyses treat cells as independent snapshots and ignore their phylogenetic relationships. Here we present LaVOUS, a lineage-aware probabilistic framework for modeling sparse single-cell gene-expression counts on reconstructed lineage trees. LaVOUS couples Brownian motion and Ornstein–Uhlenbeck models of latent transcriptional dynamics with negative-binomial observation models and scalable variational inference, enabling likelihood-based tests for gene-expression heritability, branch-specific expression shifts, and ancestral expression reconstruction. In simulations, LaVOUS improved detection of lineage-associated expression changes over Gaussian phylogenetic models and accurately reconstructed expression histories across expression levels. Applied to lineage-resolved single-cell datasets from metastatic lung cancer, class-switching B cells, and the developing brain, LaVOUS identified expression changes associated with metastatic progression, isotype switching, and neuronal differentiation. LaVOUS provides a general framework for studying single-cell expression dynamics across development and disease.

## Introduction

Understanding the precise history of cellular decision-making is fundamental to addressing major challenges in biology, from deciphering the origins of chemoresistance in cancer metastasis to mapping cell fate in organ development. While single-cell transcriptomics can reveal cell states at high resolution, it captures only a static snapshot, often obscuring the historical context of dynamic cellular changes. Recent advances in lineage tracing—especially based on the CRISPR/Cas9 barcoding technology, where heritable edits to barcodes are introduced continuously during cell growth *in vivo* [1]—now enable reconstruction of cell-lineage phylogenies at high resolution [2–7]. In some designs, CRISPR barcodes are transcribed, yielding simultaneous readouts of lineage trees and gene-expression profiles [8–10]. Unlike pseudotime trajectories inferred from transcriptomes alone [11–13], these experiments provide an independent scaffold for interpreting changes in gene expression, consisting of an explicit topology and branch lengths that reflect the true history of cell division. The resulting data sets offer a unique opportunity to study cell state changes along specific lineages, with the potential for new insights into cancer evolution, development, and cell differentiation [14–18].

Following recent progress in lineage tracing, various computational methods have been developed to reconstruct high-resolution cell-lineage trees from CRISPR barcodes [19–22]. Integrating these phylogenies with gene-expression data, however, remains challenging. Several recent methods combine lineage tracing with single-cell transcriptomics [23, 24], including approaches for lineage-aware trajectory inference [25], RNA velocity estimation [26], fate-bias identification [27], reconstruction of unobserved ancestral cell states [28, 29], quantification of transcriptional heritability and plasticity [30], and detection of lineage-associated gene-expression programs [31, 32]. These approaches are beginning to associate endpoint transcriptional states with clonal histories, but they do not yet provide explicit descriptions of how gene expression evolves along a cell-lineage phylogeny.

Evolutionary stochastic processes provide a natural framework for addressing this problem by treating gene expression as a dynamic trait evolving along the tree. These models originate from the phylogenetic comparative literature, where they have been widely used to model continuous-trait evolution, stabilizing selection, and adaptive regime shifts across species [33–35]. They have also been applied to cross-species gene-expression evolution to study regulatory divergence, drift, stabilizing selection, and lineage- or organ-specific expression evolution [36–39]. In particular, tree-based models of Brownian motion (BM) capture the accumulation of random transcriptional fluctuations, whereas the Ornstein–Uhlenbeck (OU) extension includes mean reversion toward an optimal expression level, enabling modeling of stabilizing selection. Furthermore, by allowing this optimum to vary across labeled branches or regimes, OU models can represent repeated lineage-associated shifts in these optima during processes such as tissue migration, cell differentiation, or development.

Recent methods have begun to apply such stochastic models to paired lineage-tracing and single-cell RNA-seq data. For example, EvoGeneX integrates lineage-based BM/OU dynamics with Gaussian observation models at the leaves of the tree [40]. Related approaches have been applied to single-cell cancer clone data to identify expression changes associated with aggressive or resistant phenotypes [41]. More recently, SCOUT used phylogenetic smoothing to analyze single-cell lineage trees based on expression data that had been preprocessed to better obey Gaussian assumptions [42]. These methods, however, have generally avoided directly modeling high-throughput scRNA-seq read counts, which tend to be sparse, skewed, and overdispersed, requiring flexible discrete densities such as the negative-binomial distribution [43]. Ideally, a full generative framework for the cellular evolution of gene expression would combine lineage-based stochastic dynamics with an explicit observation model for sparse single-cell read counts (see ref. [28] for an early attempt at such a model based on scVI [44]).

To address these limitations, we introduce LaVOUS (Lineage-aware Variational Ornstein–Uhlenbeck Single-cell RNA-seq analysis), a probabilistic framework for modeling single-cell gene expression dynamics on lineage trees. LaVOUS couples tree-based BM/OU latent dynamics with a negative-binomial observation model and scalable variational inference, building on the broader use of variational methods for probabilistic inference from lineage-structured single-cell data [28]. In addition, LaVOUS supports likelihood-based tests for lineage-associated expression using Pagel’s *λ* [45–47], detection of branch- or state-specific expression shifts through OU regime models, and reconstruction of gene-specific latent expression histories using Gaussian belief propagation. Together, these components provide a powerful framework for analyzing how expression changes arise along cell-lineage phylogenies. Across a variety of simulated datasets, LaVOUS improved detection of lineage-associated expression changes compared with a lineage-aware Gaussian method and reconstructed accurate latent expression histories along lineage phylogenies. We then applied LaVOUS to paired lineage and transcriptomic datasets spanning three distinct biological settings: metastatic lung cancer, where tissue migration provides branch-level state labels; B-cell class switching, where immune lineages capture immunoglobulin isotype transitions; and mouse brain development, where lineage trees record differentiation toward neuronal fates. Across these datasets, LaVOUS identified lineage-associated expression changes involved in disease progression and cell differentiation.

## Results

### Overview of the LaVOUS framework

Modeling single-cell gene expression along lineage trees involves three key challenges: defining expression dynamics on a pre-inferred tree, linking latent expression states to sparse scRNA-seq read counts, and supporting hypothesis tests or statistical inferences that address relevant biological questions.

To model gene expression on lineage trees, we follow other recent studies in leveraging the Brownian Motion (BM) and Ornstein–Uhlenbeck (OU) stochastic process, both of which are now well established in evolutionary biology [38–40]. In these models, gene expression is modeled with a multivariate Gaussian (MVG) distribution, with the lineage tree encoded through the covariance matrix. This idea originates from macroevolutionary studies and has recently been adapted to single-cell settings [33, 41, 42].

Despite the power and flexibility of these models, they are poorly matched to single-cell RNA-seq data, which is typically sparse and noisy compared to bulk RNA-seq data. In particular, scRNA-seq read counts are discrete, bounded at zero, and typically overdispersed, with variance greater than the mean. As a result, they are not well described by the continuous, unbounded MVG distributions induced by the BM and OU processes. In other applications, investigators have instead made use of flexible discrete distributions for count data, such as the negative-binomial (NB) distribution, to fit the observed single-cell read counts [43]. We therefore sought to devise a framework that would combine the BM and OU processes with an NB observational model, allowing a better fit to scRNA data while maintaining the core strengths of tree-based models of gene expression.

Our new modeling framework, LaVOUS, takes as input a fixed lineage phylogeny with branch lengths, a matched scRNA-seq count matrix, and, when available, branch or cell labels describing features such as tissue location, isotype, or cell type, which broadly correspond to discrete cellular states (**Fig. 1**). For each gene, LaVOUS models an unobserved latent expression value evolving along the tree according to the BM or OU processes in the usual way, with an induced MVG distribution at leaf cells whose covariance reflects the phylogeny and branch lengths. Latent expression values are then mapped to positive expression rates through a softplus transformation and linked to observed read counts using an NB observation model (**Fig. 1**). Because the softplus–NB observation layer makes the marginal likelihood intractable, LaVOUS approximates the latent posterior at leaves using mean-field Gaussian variational distributions (see **Methods** for details).

**Figure 1:**
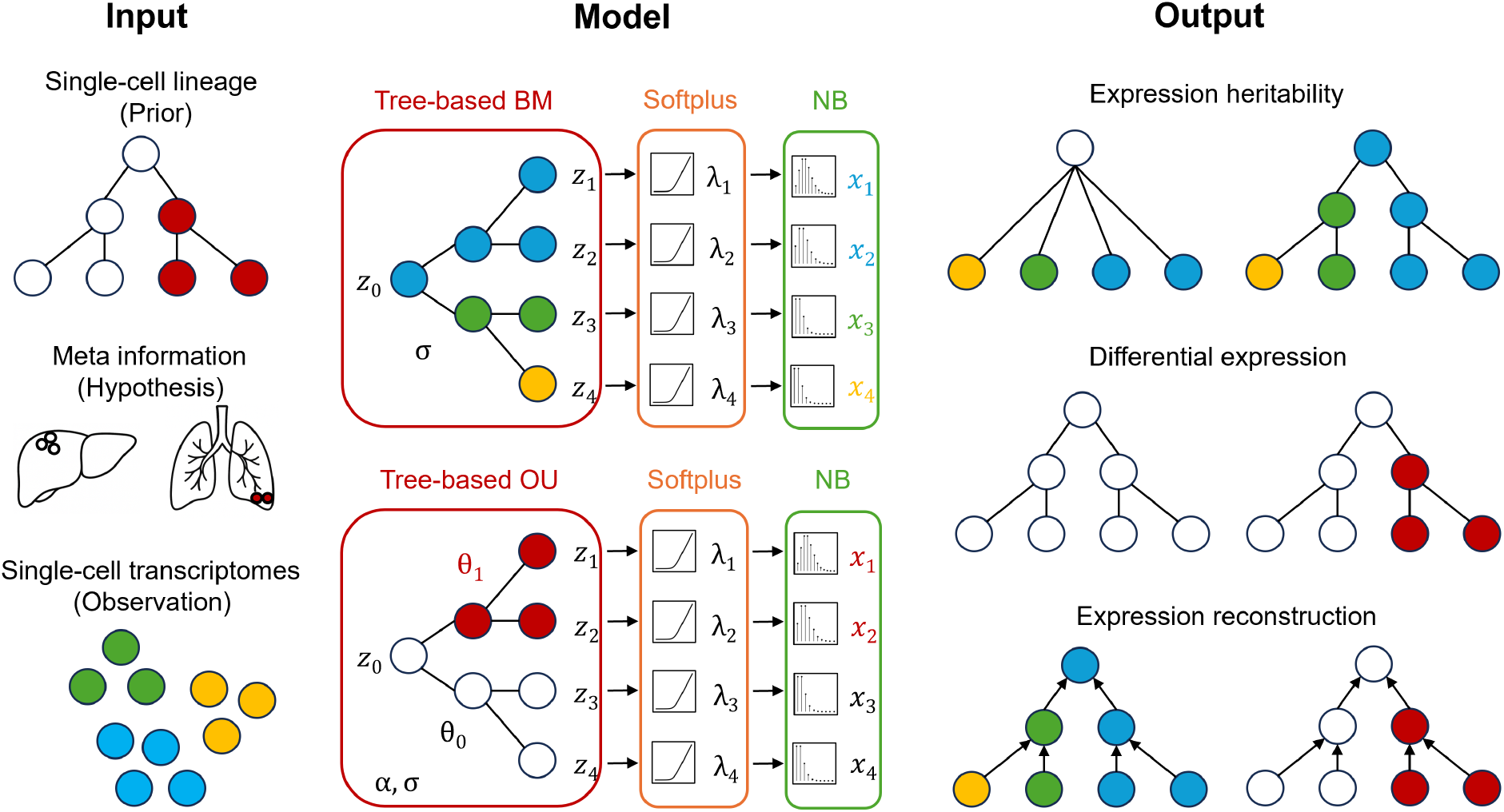
Overview of the LaVOUS framework. LaVOUS uses scRNA-seq data with a paired lineage tree as input, and identifies target genes with high heritability on the tree, or differential expression along pre-labeled branches. In addition, LaVOUS can reconstruct gene-specific latent expression histories, highlighting the putative time points of expression shifts. BM, Brownian Motion, OU, Ornstein–Uhlenbeck process. *θ*_0_, optimal expression of primary cells (e.g., primary tumor). *θ*_1_, optimal expression of other cells (e.g., metastatic tumor). *α*, selective strength of OU. *σ*, variance of BM or OU. *z*, latent expression levels of each cell from BM or OU. *λ*, positive-valued latent expression levels after softplus transformation. *x*, observed expression read counts modeled using negative-binomial (NB) sampling.

This modeling framework supports three main analyses (**Fig. 1**). First, LaVOUS tests whether a gene’s expression levels follow the phylogeny by introducing Pagel’s *λ* (*λ*_*P*_ ) [46], which scales the off-diagonal entries of the tree-induced covariance matrix. Comparing a model with estimated *λ*_*P*_ to a model with *λ*_*P*_ = 0 provides a likelihood-ratio test of whether related cells have more similar expression than expected under an independent-cell model, implying that expression levels are at least partially inherited from parent cells to their children. Second, when branches are assigned to biological regimes, LaVOUS tests for branch- or state-specific expression shifts by comparing an OU model with a shared optimum *θ* to one with state-specific optima. In addition, likelihood-ratio statistics are evaluated with asymptotic or empirical calibration and corrected for multiple testing across genes. This test enables detection of genes whose latent expression dynamics differ across states corresponding to features such as tissue locations, cell types, or developmental cell states. Third, for genes supported by the lineage-aware model, LaVOUS reconstructs gene-specific latent expression histories along internal branches and ancestral nodes using Gaussian belief propagation, providing a visualization of where expression changes are inferred to occur on the tree.

Together, these model components allow LaVOUS to describe gene expression as a continuous stochastic process following the cell division history, while accommodating sparse, overdispersed scRNA-seq read counts. In the following sections, we first evaluate LaVOUS on simulations where the true expression dynamics are known, and then apply it to paired lineage and transcriptomic datasets from metastatic lung cancer, B-cell class switching, and mouse brain development.

### LaVOUS detects single-cell gene expression patterns on simulated data

We first evaluate LaVOUS against simulated gene expression data, where the true lineage history is known and the degree of model misspecification can be controlled. We simulated phylogenies under an agent-based tumor growth model [48]. For all experiments below, we focused on a simulated lineage tree with 646 cells and one metastatic event (**Fig. 2A**). In addition, we performed the same analysis on a permutated tree with a different structure, and showed that the result is robust across different lineage trees (**Fig. S1**). To generate corresponding gene expression data from this phylogeny, we simulated 500 genes under each simulation setting, using tree-based BM and OU processes followed by softplus and NB. These simulations spanned different expression levels (*θ*), different variances (*σ* and *α*), and different overdispersions (*r*).

**Figure 2:**
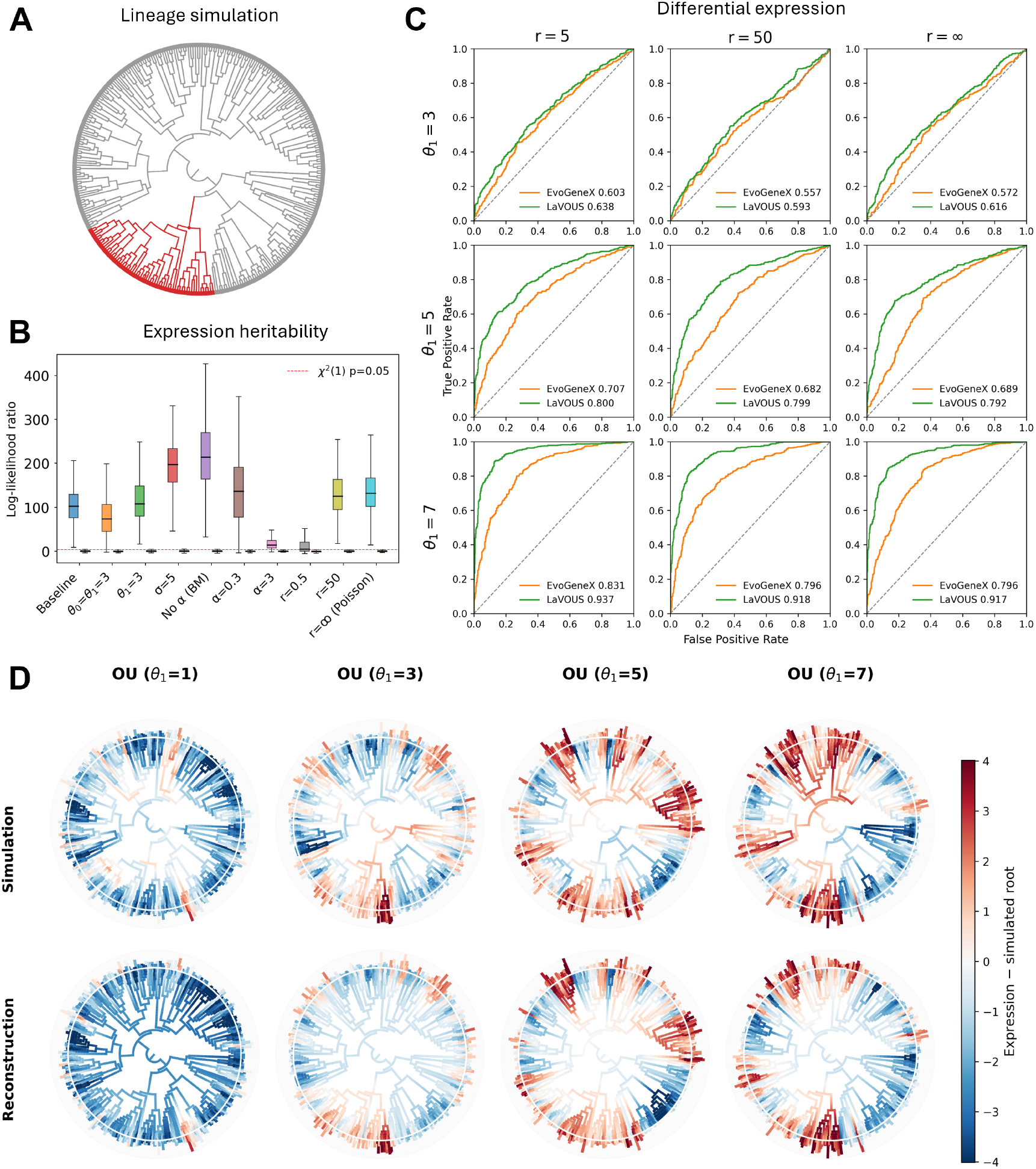
Performance of LaVOUS on simulated data. (A) Simulated cell-lineage tree from agent-based simulators in MACHINA [48]. (B) Log-likelihood ratios from heritability tests using simulated data under various settings along the *x* axis. Each setting is simulated by modifying the same baseline parameters (*α* = 1, *σ* = 3, *θ*_0_ = 1, *θ*_1_ = 1, *r* = 5). Each setting contains results from both the tree-based simulation on the left (above the threshold) and the permuted negative control on the right (below the threshold). (C) Model performance evaluated by receiver operating characteristic (ROC) curves. Rows show results for different choices of the expression optimum *θ*_1_ for the highlighted clade in (A), and columns show results for different overdispersion rates *r*. Green, LaVOUS; orange, EvoGeneX. Corresponding areas under each curve are labeled in the legend. (D) Simulations and reconstructions of gene-expression history on lineage trees under different OU models. Starting values at the root are in white. Red branches, up-regulation; blue branches, down-regulation; surrounding bars, read counts from negative-binomial sampling.

First, we applied LaVOUS to detect genes with heritable expression patterns according to the lineage tree. To compare the heritable genes with highly plastic genes whose expression levels do not follow the tree structure, we used the tree-based simulations as positive cases, and randomly permuted them across cells for negative controls. To identify heritable gene expression patterns that follow the tree structure, LaVOUS performs likelihood-ratio tests based on the BM process with free Pagel’s *λ* against Pagel’s *λ* = 0. We found that LaVOUS correctly identified genes as showing heritable gene expression from most of the positive simulations, except when the variance was exceptionally small (e.g., large *α* led to collapsed covariance) or overdispersion was exceptionally high (e.g., large *r*^−1^ led to nearly independent samples). The negative controls all correctly resulted in log-likelihood ratios close to zero (**Fig. 2B**).

To test whether gene expression levels differed significantly along selected branches, we simulated a grid of expression data spanning different expression levels and overdispersion rates on the same simulated tree. In these simulations, the background *θ*_0_ was fixed at 1, while the selected branch was assigned a different mean *θ*_1_, with larger values of *θ*_1_ indicating stronger expression up-regulation. Furthermore, larger values *r* indicated a smaller overdispersion *r*^−1^, with *r* = *∞* indicating a Poisson distribution (no overdispersion). For each simulation setting, genes simulated with different *θ*_1_ ≠ *θ*_0_ at selected branches formed the positive datasets, and genes simulated with the same *θ*_1_ = *θ*_0_ for the entire tree served as negative controls. To identify genes with expression shift at simulated metastasis, we had LaVOUS perform likelihood-ratio tests under the OU process with estimation of different *θ*s versus a null hypothesis of a uniform *θ*. We additionally benchmarked LaVOUS against a previous OU-based lineage-aware tool, EvoGeneX, which was designed for bulk RNA-seq data and uses Gaussian observation models [40]. We found that LaVOUS consistently outperformed EvoGeneX, showing larger area under the receiver operating characteristic curve (AUROC) values in all experiments, suggesting that the NB observation model results in substantially improved detection power with scRNA-seq data (**Fig. 2C**).

Notably, the empirical null distribution of the likelihood-ratio statistic deviated from the asymptotic *χ*^2^ distribution, leading to anti-conservative *p*-values when the asymptotic approximation was used directly (**Fig. S2**). We therefore implemented an empirical-null calibration procedure, in which null distributions are estimated from simulated samples under the fitted null model, and this calibration substantially reduced false-positive discoveries (**Fig. S3**).

We further evaluated the quality of the variational approximation by comparing ELBO-based likelihood estimates with importance-sampling (IS) estimates of the marginal likelihood. Although absolute ELBO estimates can deviate from marginal likelihood estimates, these deviations were largely shared between the null and alternative models causing them to cancel in likelihood-ratio statistics (**Fig. S4**). These deviations could be further reduced by empirical-null calibration (**Fig. S3**). As a result, ELBO-based likelihood ratios were well aligned with IS-based likelihood ratios, particularly for genes near the non-significant range, supporting the practical use of ELBO estimates for likelihood-ratio testing in LaVOUS (**Fig. S4**).

Finally, we used LaVOUS to reconstruct gene-expression histories on lineage trees. Using our tree-based OU simulations as example data, we reconstructed these histories according to the OU and variational parameters estimated from the model (**Fig. 2D**). Although these parameters were not assigned direct biological interpretations, they specify the latent expression dynamics that enable reconstruction of gene-expression histories along lineage trees (**Fig. S5**). In particular, LaVOUS uses the estimated variational parameters as initial Gaussian parameters at the leaves of the tree and the OU model parameters for Gaussian belief propagation, producing reconstructed expression histories at each node of the tree (see **Methods**). We found that these reconstructions generally agreed well with the ground truth from our simulations (**Fig. S6**).

### LaVOUS reveals dynamic gene expression in lung-cancer metastasis

For our first real-data application, we considered a comprehensive dataset describing metastatic lung cancer in a mouse model [17]. This study provides scRNA-seq paired with high-resolution lineage trees derived from CRISPR barcodes. We focused our analysis on the largest metastatic clone from this study, which contains 20,992 cells from three tissues: lung (primary tumor), liver (metastatic tumor), and soft tissue (metastatic tumor). Many of these cells had redundant barcodes, however, so we reduced the data to 1,453 uniquely barcoded cells. We reconstructed a cell-lineage tree for these cells using VINE [22] (**Fig. 3A**), and then removed genes present in fewer than 10% of cells, leaving nearly eight thousand genes for downstream analysis. To reduce technical biases from differences in sequencing depth, we additionally scaled the read counts for each cell by the corresponding median library size (see **Methods**).

**Figure 3:**
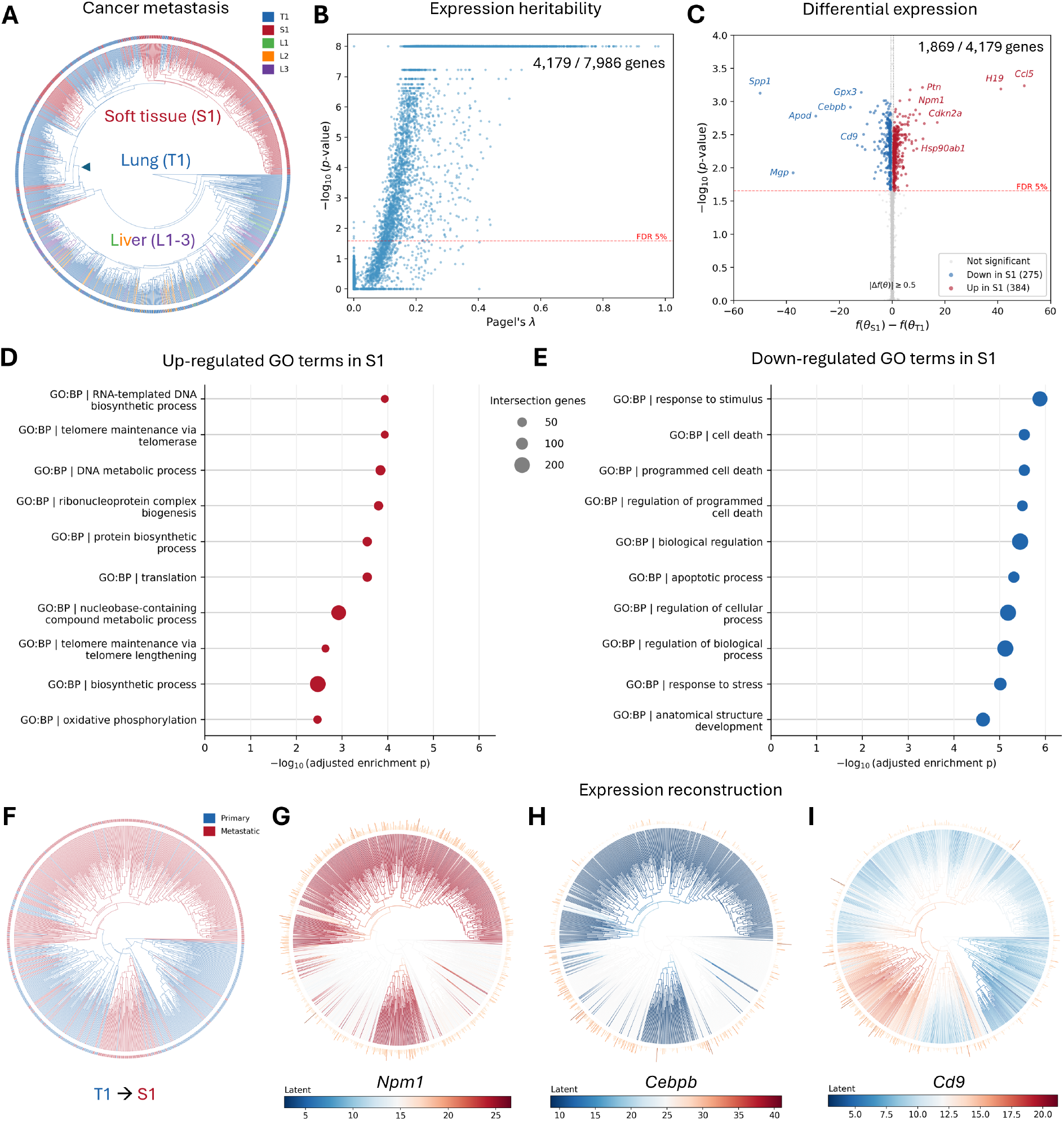
Application of LaVOUS to scRNA-seq data describing metastatic lung cancer in a mouse model [17]. (A) Reconstructed lineage tree colored by tissue label. Arrow shows the root of the clade considered in tests of differential expression. T1, primary tumor in lung; S1, metastatic tumor in soft tissue; L1–3, metastatic tumor clones in liver. (B) Heritability test applied to all genes. (C) Differential-expression test applied to heritable genes, considering all the T1-to-S1 branches in the clade of interest. Genes with absolute effect sizes equal to or larger than 0.5 (difference of *θ* after softplus on the *x* axis) and adjusted *p <* 0.05 after empirical-null calibration are considered significant targets. (D) Gene Ontology (GO) enrichment analysis for significantly up-regulated genes. (E) GO enrichment analysis for significantly down-regulated genes. (F) The full T1-to-S1 clade, colored by tissue. (G–I) Gene expression histories reconstructed for three representative genes. Root is colored in white, and up/down regulation is indicated in red/blue, respectively.

Out of 7,986 genes, LaVOUS identified slightly more than half as having significant evidence of gene-expression heritability along the tree (FDR *<* 5%), and we focused on these genes in downstream lineage-based analyses (**Fig. 3B**). Since the reconstructed tree indicates a complex, multi-tissue history of metastasis, we focused our analysis of differential expression on one large clade in which only the primary tumor (T1) and one secondary tissue (soft tissue S1) were represented (**Fig. 3A, F**). In a test of differential expression on all T1*→*S1 branches, 1,869 genes showed significant up- or down-regulation in S1 after empirical calibration (see **Methods**), including 384 up-regulated genes and 275 down-regulated genes with mean difference ≥ 0.5 (**Fig. 3C**). The up-regulated genes were enriched for roles in migration, colonization, and proliferation (**Fig. 3D**). For example, *H19* is an lncRNA facilitating metastasis [49], *Ptn* is a growth factor for angiogenesis and tumorigenesis that may modify the microenvironment [50], and *Npm1* is a nucleolar protein involved in ribosome biogenesis [51]. In contrast, down-regulated genes such as *Apod, Cebpb*, and *Gpx3* are enriched for roles in stress response, suggesting adaptation to the new microenvironment [52–54] (**Fig. 3E**).

In addition, reconstruction of gene-expression levels along the cell lineage tree revealed distinct patterns for different gene targets. For instance, genes such as *Npm1* and *Cebpb* are strongly up- or down-regulated in the metastatic tumor, with the shift in gene expression coinciding with the metastatic event (**Fig. 3G, H**). In contrast, *Cd9*, which encodes a cell surface glycoprotein for cell adhesion, has complex expression history in metastatic and primary branches, consistent with its context-dependent functions in migration, colonization, and proliferation [55] (**Fig. 3I**). Taken together, these results show that LaVOUS can both detect novel gene targets and illuminate gene-expression dynamics through lineage-aware reconstruction.

### LaVOUS characterizes isotype switching in B-cell immune response

Next we applied LaVOUS to a dataset describing the gene-expression response of human B cells to the SARS-CoV-2 mRNA-1273 vaccine [56]. In this case, B-cell lineage trees can be inferred from natural markers such as somatic hypermutations (SHM) during antibody generation. We collected preprocessed data from Weber et al. [57] and reconstructed B-cell lineages using the IgPhyML method from the Dowser package [58]. The isotype labels on the trees were then inferred using Fitch parsimony.

In our analysis, we focused on the two dominant immunoglobulin isotypes in the dataset: IgG1 and IgA1. IgG1 is commonly induced by intramuscular vaccine injections and is responsible for blood or systemic immunity [59]. Under class-switching recombination (CSR), IgG1 can switch to IgA1, which is more effective for airway or mucosal immunity [60, 61] (**Fig. 4A**). Here, we used the two largest clonotypes with clear IgG1-to-IgA1 transitions for downstream gene expression analyses (**Fig. 4B**). Because individual B-cell lineage trees are relatively small, we increased statistical power by jointly analyzing two independent clonotypes (by summing their log likelihoods). Our analysis identified 49 genes with heritable gene expression, of which 37 genes are differentially expressed in the IgA1 isotype. Further calibration using gene-specific empirical-null distribution reduced the number of significant genes to ten (with 10% FDR). Specifically, *XBP1*, a gene associated with active antibody secretion in plasmablast B cells, is significantly down-regulated in IgA1 branches [62] (**Fig. 4C, D**), suggesting that the newly formed IgA1 B cells may be still in premature states in transit to tissue relocation, instead of in plasmablast states with active antibody secretion. In contrast, *JCHAIN*, another gene typically used for antibody joining in IgA, is up-regulated in IgA1 branches (**Fig. 4E, F**). Antibody dimerization through this mechanism helps epithelial transport of IgA in mucosal tissues, which is especially critical for respiratory infections such as COVID-19 [63]. Taken together, these results demonstrate that LaVOUS enables characterizations of B-cell isotype transitions during immune responses, and provides explicit expression histories for highlighted gene targets in B-cell development.

**Figure 4:**
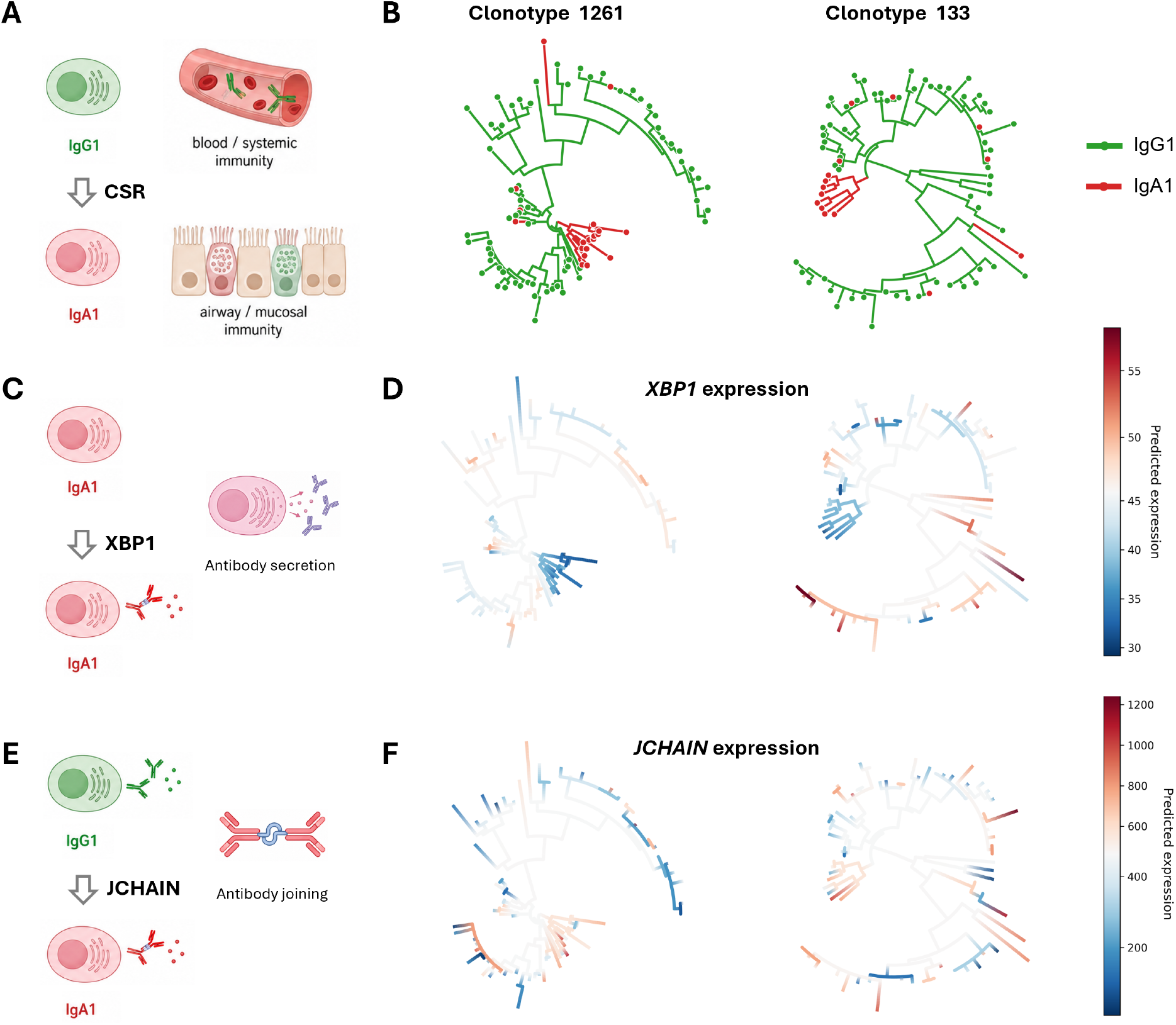
Application of LaVOUS to scRNA-seq data for human B cells treated with SARS-CoV-2 mRNA-1273 vaccine [56]. (A) Isotype transition from IgG1 B-cell to IgA1 B-cell through class-switching recombination (CSR). (B) B-cell lineage trees from two clonotypes. Isotypes are labeled in different colors. (C) Biological function of *XBP1* gene. (D) Gene expression reconstruction for *XBP1* on B-cell lineages. Predicted latent expressions are labeled in different colors. The root values are in white, and the red or blue branches indicate up- or down-regulation, respectively. (E) Biological function of *JCHAIN* gene. (F) Gene expression reconstruction of *JCHAIN* on B-cell lineages.

### LaVOUS highlights developmental signals in ventral midbrain of mouse

Finally, we applied LaVOUS to scRNA-seq data for the developing ventral midbrain of the mouse [64]. Previous work identified differentiation paths for dopaminergic neurons (DA) and glutamatergic neurons (GLU) from radial glia-like cells (Rgl) and neuroblasts (Nb) in the floor plate (FP) region [64] (**Fig. 5A**).

**Figure 5:**
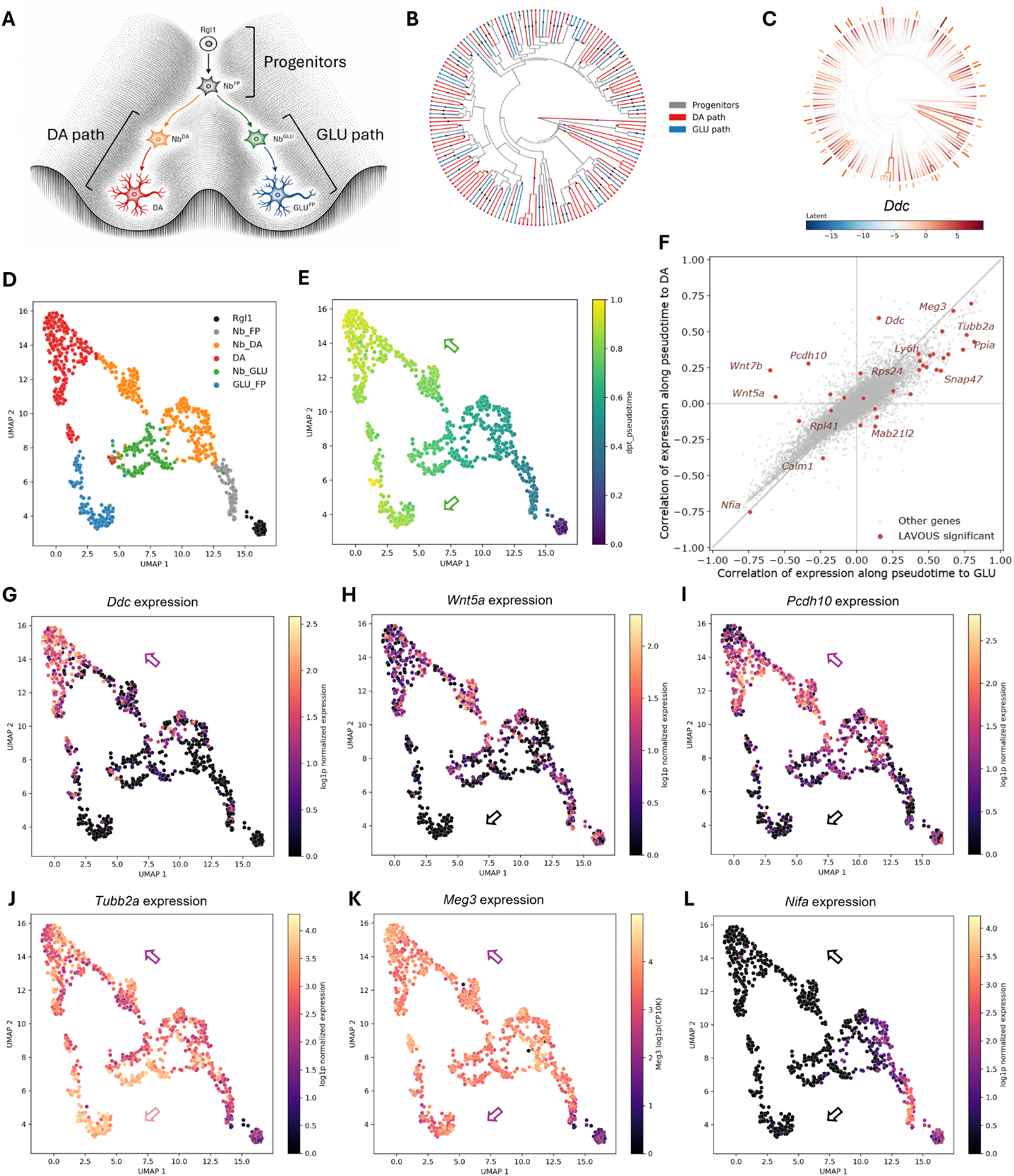
Application of LaVOUS to data for mouse brain development [64]. (A) Differentiation from progenitors to neurons at floor plate (FP) of mouse ventral midbrain. Rgl, radial glia-like cell; Nb, neuroblast; DA, dopaminergic neuron; GLU, glutamatergic neuron. (B) Lineage tree of FP clones based on CRISPR barcodes. Cell types are labeled by color. Differentiation events are highlighted with dots on branches. (C) Gene expression reconstruction of *Ddc* gene. (D) UMAP of all FP cell transcriptomes colored by cell types [69]. (E) Same UMAP colored by diffusion-pseudotime trajectory [12]. Arrows show developmental paths to DA and GLU. (F) Pearson correlations in gene expression changes along the pseudotime trajectories in the DA path (*y* axis) and GLU path (*x* axis). Significant genes identified by LaVOUS are highlighted in red. (G–L) Same UMAP colored by *Ddc, Wnt5a, Pcdh10, Tubb2a, Meg3*, and *Nfia* expression, respectively. Purple arrows show positive correlations (up-regulation) along the pseudotime trajectory; black arrows show negative correlations (down-regulation).

We selected clones with FP cell types from the E15-rep1 sample and reconstructed the lineage tree by VINE using the mutation patterns in the available CRISPR barcodes [22] (**Fig. 5B**). We also reconstructed the ancestral cell types on the tree by following the differentiation paths from progenitors to neurons. To identify genes with differential expression in the DA- and GLU-differentiating paths, we compared an alternative hypothesis of three *θ*s for progenitors, DA-path cells, and GLU-path cells, respectively, with a null hypothesis of two *θ*s for progenitors and DA/GLU-path cells (**Fig. 5A**). As a result, LaVOUS identified 68 genes with heritable expression based on all the clones, of which 35 genes are significantly different in DA-path and GLU-path cells after empirical-null calibration based on the FP clones. For example, *Ddc*, which encodes aromatic L-amino acid decarboxylase involved in dopamine and serotonin synthesis, was significantly up-regulated in DA cells [65] (**Fig. 5C**).

To assess how LaVOUS complements transcriptome-based analyses, we compared lineage relationships with UMAP embedding and pseudotime trajectories. Although cells of the same type often have different origins in the lineage tree, they tend to group in a UMAP visualization of the transcriptomes, suggesting that gene expression data are insufficient and barcoding is needed for accurate lineage-tree reconstruction (**Fig. 5D**). For comparison with our lineage-based analysis, we performed trajectory inference using Scanpy and identified pseudotime evolution from Rgl1 to DA and GLU, confirming the two differentiation paths previously reported [64, 66] (**Fig. 5E**). To identify differentially expressed genes during the development of DA and GLU along the pseudotime trajectory, we compared correlations of gene expression changes with the pseudotime changes in the two developmental paths, highlighting significant genes from LaVOUS as a reference (**Fig. 5F**). We found that some genes identified by LaVOUS agreed well with the pseudotime trajectory but others did not. For example, expression of the DA marker *Ddc* is positively correlated with pseudotime in the DA path but weakly correlated in the GLU path, indicating up-regulation specifically in DA and NbDA (**Fig. 5G**). By contrast, expression of a developmental signal *Wnt5a* is negatively correlated with the GLU path but not the DA path, indicating down-regulation specific to GLU and NbGLU [67] (**Fig. 5H**). Expression of *Pcdh10*, which encodes a cadherin that contributes to cell-cell connections in the brain, is positively correlated with DA but negatively correlated with GLU, indicating opposite expression regulations in neuron differentiation [68] (**Fig. 5I**).

At the same time, the expression levels of some other significant genes from LaVOUS showed less obvious changes along the pseudotime trajectories. Specifically, the tubulin gene *Tubb2a* is positively correlated with both the DA path and the GLU path, with only a slightly lower correlation with the latter (**Fig. 5J**). Likewise, the lncRNA *Meg3* has similar positive correlations with both DA and GLU, and the transcription factor-encoding gene *Nfia* is similarly negatively correlated in both paths (**Fig. 5K–L**). These observations suggest that LaVOUS sometimes has improved sensitivity for dynamic gene expression by considering true cell lineages rather than relying on the more approximate pseudotime trajectory. Overall, LaVOUS reveals dynamic biomarkers and developmental signals during mouse brain development, providing insights beyond pseudotime trajectories.

## Discussion

In this study, we have introduced LaVOUS, a modeling framework for analyzing the dynamics of gene expression along a given lineage phylogeny. LaVOUS takes reconstructed cell-lineage trees and paired scRNA-seq data as inputs, models gene expression evolution on the tree as a stochastic process, and computes the likelihood of sparse, overdispersed read counts using negative-binomial observation models. LaVOUS supports several types of analysis, including the evaluation of gene-expression heritability using Pagel’s *λ*, differential gene expression using OU processes, and evolutionary reconstruction using Gaussian belief propagation. We validated LaVOUS extensively on simulated data, and showed that its incorporation of negative-binomial observation models with variational inference, in particular, substantially improves statistical power for gene-target identification.

To demonstrate the broader utility of LaVOUS, we additionally applied it to three representative datasets ranging from lung-cancer metastasis to B-cell immune responses and neural development, revealing aspects of the dynamics of disease progression and cell differentiation. Notably, these datasets not only spanned diverse biological processes, but they also involved a range of model applications and consideration of markedly different tree structures. For instance, the cancer cells have high proliferation rates, resulting in large clonal trees with clear shifts among tissue labels. In this setting, LaVOUS was able to identify large enough numbers of gene targets to support GO-enrichment analysis and visualization of complex patterns of gene expression along the phylogeny. In contrast, the B-cell lineages decompose by clonotype, which leads to smaller lineage trees. As a result, we used LaVOUS to jointly infer likelihoods across multiple trees, enabling identification of more gene targets by pooling information from multiple clonotypes. Finally, the developmental dataset contains cells from a variety of developmental stages and highly interspersed cell type labels on the tree, because differentiated neurons cease to divide. As a result, LaVOUS identifies developmental signals for dopaminergic and glutamatergic neurons, many of which are not present in pseudotime trajectories that are not faithful to the true lineage history.

In the case of the lung-cancer data sets, we were particularly interested to find multiple lines of evidence suggesting that metastatic cells in soft tissue are actively proliferating and undergoing clonal expansions. First, biosynthetic genes such as *Npm1* are up-regulated, and stress-response genes such as *Cebpb* and *Gpx3* are down-regulated, consistent with enhanced proliferative or biosynthetic activity (**Fig. 3C**). Second, GO-enrichment analysis of differentially expressed genes identified biosynthesis and metabolism as upregulated terms, and death, apoptosis, and stress-response as down-regulated terms, suggesting more active cell states in the metastatic tumor (**Fig. 3D, E**). Finally, the reconstructed tree indicates that metastatic cells in soft tissue emerge rapidly, early in the evolutionary process (**Fig. 3A, F**). Together, these observations suggest rapid colonization of the soft tissue owing to a high proliferation rate and/or reduced death following migration from the primary tissue.

LaVOUS could benefit from several extensions to the current model. For example, the OU process currently assumes constant evolutionary rates along all branches of the lineage tree, whereas the true process likely obeys different rates at different time points, owing to intrinsic heterogeneity or changing environmental signals. It is straightforward to accommodate such rate heterogeneity in the model, but it remains to be seen to what extent differences in rate can be accurately inferred from data. Another useful extension would be to consider paired time-series data, which could help to address the missing information on the lineage tree. The performance of LaVOUS naturally depends on the provided tree structure, which can be difficult to reconstruct accurately with current data. This dependency could be mitigated by adapting LaVOUS to integrate over samples from the posterior distribution of trees, as enabled by recent Bayesian methods [21, 22]. Finally, LaVOUS models the dynamics of gene expression independently at each gene, whereas in reality genes are strongly correlated. A longer-term goal would be to extend the model to consider collections of genes together, to provide insights into single-cell dynamics at the level of regulatory networks.

In summary, LaVOUS combines single-cell data describing both transcriptomes and lineage tracing, and enables a rich collection of analyses of gene expression dynamics along a phylogeny. We anticipate that LaVOUS will help to make lineage-aware modeling a central component of future single-cell analyses, particularly when combined with spatial, multi-omic, and perturbational measurements.

## Materials & Methods

### Model for stochastic gene expression along cell-lineage trees

We model the latent expression of each gene along a cell-lineage tree using either Brownian motion (BM) or the Ornstein–Uhlenbeck (OU) process [37–42]. Under BM, the latent expression state *z*_*t*_ evolves according to the stochastic differential equation (SDE),

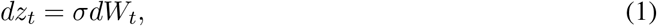

where *σ >* 0 is the diffusion coefficient and *W*_*t*_ is a standard Wiener process. Over a branch of length *t* from cell *i* to cell *i* + 1, this SDE induces a Gaussian transition distribution,

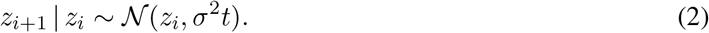

Similarly, under the OU process,

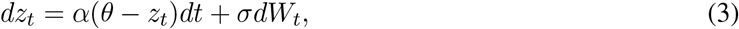

which induces the transition distribution,

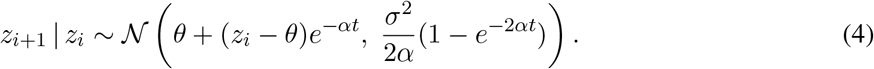

In this case, *α* controls the strength of attraction toward an optimal expression level *θ*.

Using Eq. 2 or Eq. 4, we model the latent gene expression *z* along each branch of the lineage tree *T* from the root to the leaves. For BM, we start with a mean expression *µ*_0_ as a free parameter at the root. As a result, the joint gene expression levels at the leaves of the tree, denoted ***z***, follows a multivariate Gaussian (MVG) distribution with likelihood,

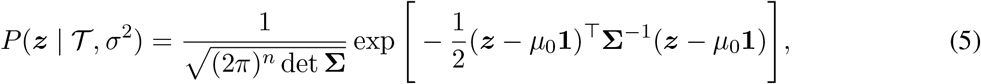

where **Σ** is the covariance matrix. The covariance between leaves *i* and *j* is given by Σ_*ij*_ = *σ*^2^*s*_*ij*_, where *s*_*ij*_ is their shared branch length in *T* from the root to their most recent common ancestor (MRCA).

For the OU process, we assume at the root of the tree that *z*_0_ is at stationarity for the initial regime, represented by a Gaussian prior 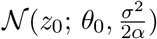. As a result, the joint gene expression levels at the leaves follow an MVG with likelihood,

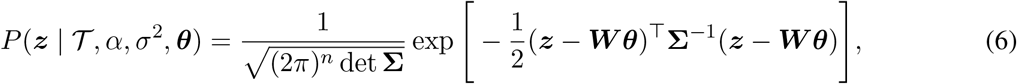

where the mean components are defined by weighted sums of ***θ*** elements according to a weight matrix ***W*** such that,

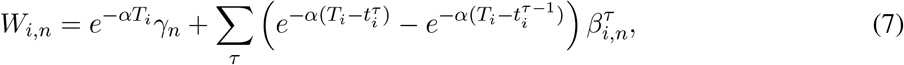

where *T*_*i*_ is the total branch-length from leaf *i* to the root, 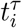 is the total branch-length from the root to the terminal point of regime *τ* along the lineage leading to leaf *i, γ*_*n*_ is a binary indicator of whether *θ*_*n*_ is active at the root, and 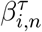 is a binary indicator of whether *θ*_*n*_ is active for cell *i* in regime *τ*, as detailed in ref. [40]. To define the expression regimes for this model, cell labels for internal nodes are either inferred using VINE (with CRISPR barcodes) or by Fitch parsimony (otherwise) [19, 22]. Under this model, the covariance between leaves *i* and *j* depends on both the total branch-lengths from the root to the leaves (*T*_*i*_ and *T*_*j*_) and the branch lengths from the root to the MRCA (*s*_*ij*_):

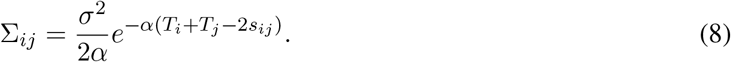

In practice, we optimize the MVG likelihood subject to the Karush-Kuhn-Tucker (KKT) constraints, as described in ref. [40].

### Observation model for single-cell read counts

The BM and OU processes provide convenient MVG likelihoods for continuous expression levels given the tree structure. Single-cell read counts, however, are discrete, nonnegative, and overdispersed proxies for expression levels, which are not well matched to Gaussian assumptions [43]. We therefore introduce an observation model based on a softplus transformation and negative-binomial (NB) distribution (see also ref. [28]). Specifically, each latent Gaussian expression level *z*_*i*_ from the tree-based process is first transformed to *λ*_*i*_ *∈* (0, *∞*) by the softplus function,

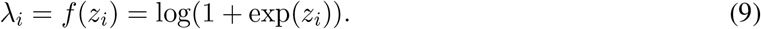

The read counts are then described using an NB distribution with mean *λ*_*i*_ and dispersion parameter *r*, such that the variance is given by 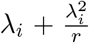 (meaning that small *r* implies overdispersion). For simplicity, we assume a uniform *r* across all leaves, resulting in the following conditional likelihood for read counts ***x*** given transformed latent expression levels ***λ***,

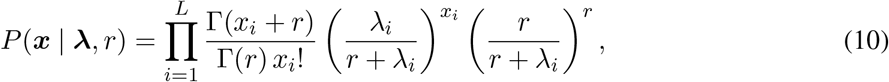

where *L* is the number of leaves of the tree.

Notably, if cell-specific library size scaling factors *l*_*i*_ are available (such that *l*_*i*_ normalizes cell *i* relative to the median value for all cells), we replace *λ*_*i*_ with *l*_*i*_*λ*_*i*_ to accommodate library-size differences.

### Approximation for full likelihood based on variational inference

The full likelihood function for observed gene expression read counts is given by,

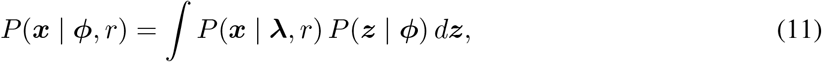

where ***ϕ*** = (*T, α, σ*, ***θ***) summarizes the tree and all BM/OU parameters. This high-dimensional integral, however, has no closed-form solution and is not amenable to standard Monte Carlo approximations. Therefore, following ref. [28], we applied mean-field variational inference to approximate the posterior distribution for the expression levels ***z*** using a product of Gaussian distributions,

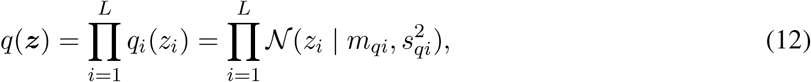

where ***m***_*q*_ and ***s***_*q*_ are chosen to minimize the Kullback–Leibler (KL) divergence to the true posterior,

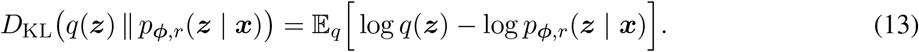

By replacing the true posterior using Bayes’ theorem,

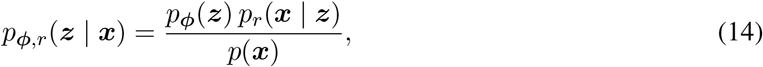

we obtain a tractable evidence lower bound (ELBO) for variational inference,

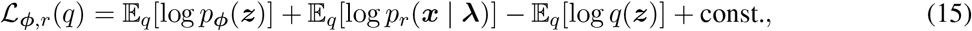

which serves as a strict lower bound for the log likelihood. The first term has a closed-form expression. In the case of BM,

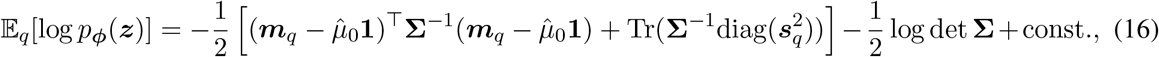

and for OU,

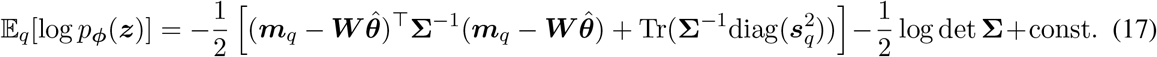

The second term, representing the softplus–NB observation model, reduces to,

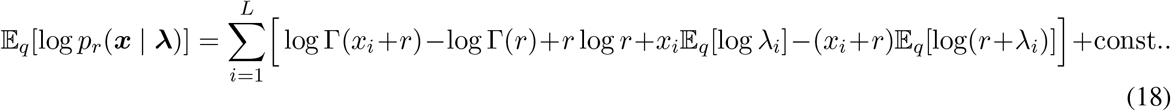

The remaining expectations were estimated using Monte Carlo samples with reparameterization. For the last term, the entropy,

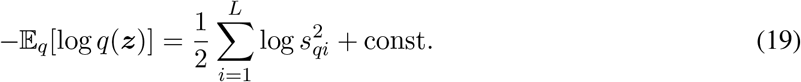

We jointly estimate the model and variational parameters by numerically maximizing Eq. 15 using the Adam optimizer in PyTorch. Notably, the trailing constant does not depend on the model parameters and can be ignored during optimization.

In cases where it is useful to pool data across trees, we sometimes optimize a joint log-likelihood function (sum of log likelihoods) based on shared parameters.

### Likelihood-ratio test to detect gene expression heritability along the tree

To test whether the expression of a gene is heritable along the cell-lineage tree, we made use of Pagel’s *λ* (here denoted *λ*_*P*_ ), a scalar parameter that controls the strength of phylogenetic correlation among cells [46]. Given the covariance matrix **Σ** defined by the input lineage tree, we construct a *λ*_*P*_ -transformed covariance matrix,

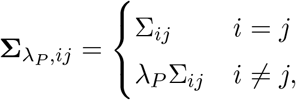

where *λ*_*P*_ *∈* [0, 1]. When *λ*_*P*_ = 1, the full tree-induced covariance is retained, corresponding to a BM model in which related cells have correlated latent expression values. When *λ*_*P*_ = 0, all off-diagonal covariance terms are removed, corresponding to a star-tree or independent-cell model in which shared lineage histories make no contribution to patterns of expression variation. Intermediate values of *λ*_*P*_ allow for partial contributions of heritability between cells.

For each gene, we compare two models. Under the null hypothesis *H*_0_, we fix *λ*_*P*_ = 0, so that the latent expression values are independent across cells. Under the alternative hypothesis *H*_1_, *λ*_*P*_ is estimated jointly with the remaining model parameters, allowing the data to determine the extent to which expression variation follows the tree. In both models, latent expression values evolve according to the *λ*_*P*_ -weighted BM and are linked to observed read counts through the softplus–NB observation model described above.

The variational likelihood-ratio test statistic is approximated by the difference between the ELBOs,

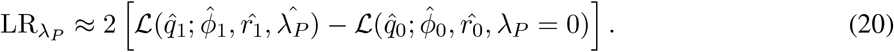

Because the alternative model introduces one additional free parameter, *λ*_*P*_, we evaluate significance approximately using a *χ*^2^ distribution with one degree of freedom. Genes with significant likelihood-ratio test statistics are interpreted as showing support for lineage-associated, heritable expression patterns. Finally, we apply the Benjamini–Hochberg procedure across genes to control the false discovery rate (FDR).

### Likelihood-ratio test for branch-specific shifts in gene expression

We test for expression shifts across cell types or tissue locations using an OU-based framework [37– 40]. Under the null hypothesis *H*_0_, all cells share a single optimal expression level *θ*, whereas under the alternative hypothesis *H*_1_, each label of interest has a distinct *θ*. More generally, *H*_0_ may specify multiple *θ* values, provided its label structure is nested within that of *H*_1_.

Similar to the heritability test, we perform a variational likelihood-ratio test using ELBO approximations:

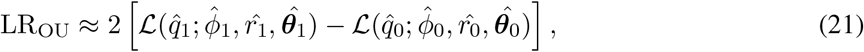

with degrees of freedom equal to the difference in the number of *θ* parameters between hypotheses. Significance is assessed against both a *χ*^2^ reference distribution and an empirical null distribution obtained by simulating under the fitted null model and re-fitting both hypotheses. The empirical null can be shared across all genes or computed per gene. Optionally, its tail is fitted with a generalized Pareto distribution (GPD) to reduce the number of simulations. FDR was controlled using the Benjamini–Hochberg procedure.

### Diagnostic assessment of the ELBO approximation

By Jensen’s inequality, the ELBO provides a lower bound on the marginal log likelihood,

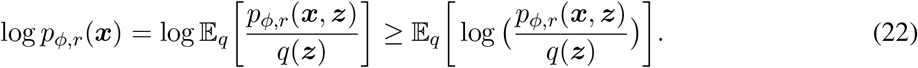

To assess the accuracy of the ELBO approximation, we also estimated the marginal log likelihood using importance sampling (IS),

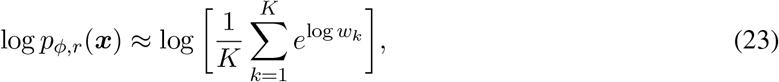

where *K* is the number of importance samples. To improve coverage of the latent space, samples were drawn from a mixed proposal distribution,

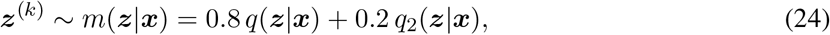

where *q*_2_(***z***|***x***) has the same mean as *q*(***z***|***x***) but twice its standard deviation. The corresponding log importance weight was,

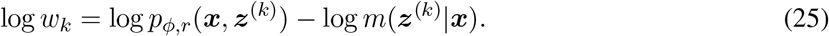

Using this IS-based estimate as a diagnostic, we found that likelihood-ratio statistics computed from ELBOs were close to those computed from IS-based marginal likelihood estimates near the significance threshold, suggesting that the ELBO approximation is adequate for likelihood-ratio testing in differential expression (Fig. S4). However, because the effective sample sizes of IS were generally small in this high-dimensional latent space, we used IS only as a diagnostic and relied on ELBO-based likelihood ratios for the main analyses.

### Gaussian belief propagation to reconstruct gene expression history along the tree

For genes identified as heritably and differentially expressed along labeled branches, we reconstruct latent gene expression histories on the full lineage tree, including internal ancestral nodes. The reconstruction is performed using Gaussian belief propagation under the fitted OU process. This procedure combines information from the variational posterior distributions at the leaves and the Gaussian transition structure imposed by the lineage tree.

In particular, after fitting the LaVOUS model for a given gene, we use the optimized variational distributions at the leaves as approximate Gaussian observations of the latent expression states:

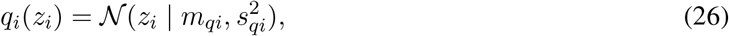

where 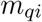 and 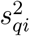 are the optimized variational mean and variance for leaf cell *i*. These variational distributions summarize the information provided by the observed read counts ***x*** after accounting for the nonlinear softplus–NB observation model.

The latent expression state evolves along each branch according to the OU process as described by Eq. 4. Because both the tree prior and the approximate leaf likelihoods are Gaussian, posterior inference over all internal and leaf latent states can be carried out efficiently by message passing. Briefly, in the upward pass, each leaf sends its Gaussian variational information to its parent. Internal nodes recursively combine Gaussian messages from their children with the branch transition densities, producing a Gaussian summary of the conditional likelihood of all descendant observations. In the downward pass, information from the root and from sibling subtrees is propagated back toward the leaves, yielding the marginal posterior distribution of each node’s latent expression state conditional on all observed cells.

For each node *v* in the tree, the result is an approximate posterior distribution,

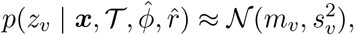

where *m*_*v*_ is the approximate posterior mean expression level and 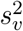 is the posterior variance. The mean *m*_*v*_ is used as the reconstructed expression history, while 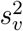 provides a measure of uncertainty in the inferred ancestral state.

### Data processing

The cancer metastasis dataset includes matched CRISPR barcodes and scRNA-seq data [17]. We focused our analysis on the largest metastatic tumor, 3724 NT All. CRISPR editing sites were used to reconstruct the sample lineage tree using VINE in migration mode [22]. The heritability test was based on the full clonal tree, while the differential test was limited to the large clade that contained only lung-to-soft tissue metastasis (**Fig. 3F**). For transcriptomes, the cells with the highest sequencing depths were selected as representatives for all cells with matching barcodes. LaVOUS was fitted to raw read counts, with median-normalized total read counts used as library-size factors. Genes that were expressed in fewer than 10% of representative cells were omitted. The differential test was applied to all genes that passed the heritability test. An empirical calibration was carried out using 20,000 simulations, with generalized Pareto distribution (GPD) tail fitting to define the empirical null distribution for all genes. Enrichment analyses of significant genes were performed with g:Profiler [70].

The human B-cell dataset includes lineages from B-cell receptors and matched scRNA-seq data [56]. Preprocessed sequence data was retrieved from the TRIBAL study [57]. Lineage trees were built by Dowser using IgPhyML with a junction length of 30 bp [58]. Our analysis focused on the two largest clonotypes, 1261 and 133, both of which exhibited clear IgG1-to-IgA1 isotype switches after removing other isotypes. Cell-type labels were inferred by Fitch parsimony, with the root state fixed to IgG1. One cell with reversed class switching was omitted. Subsequent preprocessing steps were similar to those described above. Owing to the small number of genes, however, in this case we used gene-wise 2,000 simulations as empirical null distributions.

The brain developmental dataset also includes matched CRISPR barcodes and scRNA-seq data [64]. We focused our analysis on the E15-rep1 clone, which contained sufficient numbers of floor plate cells. For the heritability test, representative cells were selected from the dominant cell type of each barcode clone. For the differential test, representative cells were selected from barcodes associated with floor-plate cell types. Cell types on trees were labeled by Sankoff parsimony with a directional matrix following the cell differentiation paths. Lineage trees were constructed by VINE and LaVOUS analyses were performed as described above. Diffusion-pseudotime (DPT) trajectory analyses were conducted using Scanpy [12, 66]. Count data were filtered for genes expressed in at least 3 cells, library-size normalized to 10,000 counts per cell, log-transformed, and reduced to the top 2,000 highly variable genes. After scaling, we computed 50 PCA components, built a 30-nearest-neighbor graph, and inferred diffusion maps and DPT. The trajectory was rooted in the Rgl1 progenitor population, choosing the root cell nearest the progenitor PCA centroid. For each gene, log-normalized expression was correlated with DPT separately along the progenitor-to-DA and progenitor-to-GLU paths, and these branch-specific Pearson correlations were compared with LaVOUS-significant genes highlighted in **Fig. 5F**.

## Code availability

LaVOUS has been implemented as an open-source Python package available under the MIT License at GitHub: https://github.com/Jiawei-Xing/LaVOUS. Generative AI tools (including ChatGPT and Claude) were used to assist with language editing, code development, and improving the clarity of the manuscript.

## Acknowledgments

We thank members of the Siepel and Nowak labs, as well as the CSHL Cancer Center and Single-Cell Core facility for support.

## Funding

This work was performed with assistance from US National Institutes of Health (NIH) National Cancer Institute (NCI) grants R01-CA272466 (to D.G.N.), NIH National Institute of General Medical Sciences grant R35-GM127070 (to A.S.), Starr Cancer Consortium grant I16-0060 (to D.G.N.), an American Cancer Society Research Scholar Grant (to D.G.N.), a Weill Cornell Medicine Walter B. Wriston Research Scholar Award (to D.G.N.), a National Science Foundation Graduate Research Fellowship (to S.J.S.), a Starr Centennial Scholarship from the Starr Foundation (to S.J.S.), a John Ligas Memorial Award (to J.X.) and the Simons Center for Quantitative Biology at CSHL. The content is solely the responsibility of the authors and does not necessarily represent the official views of the US National Institutes of Health.

## Conflicts of Interest

The authors report no conflicts of interest.

---

**Figure S1:**
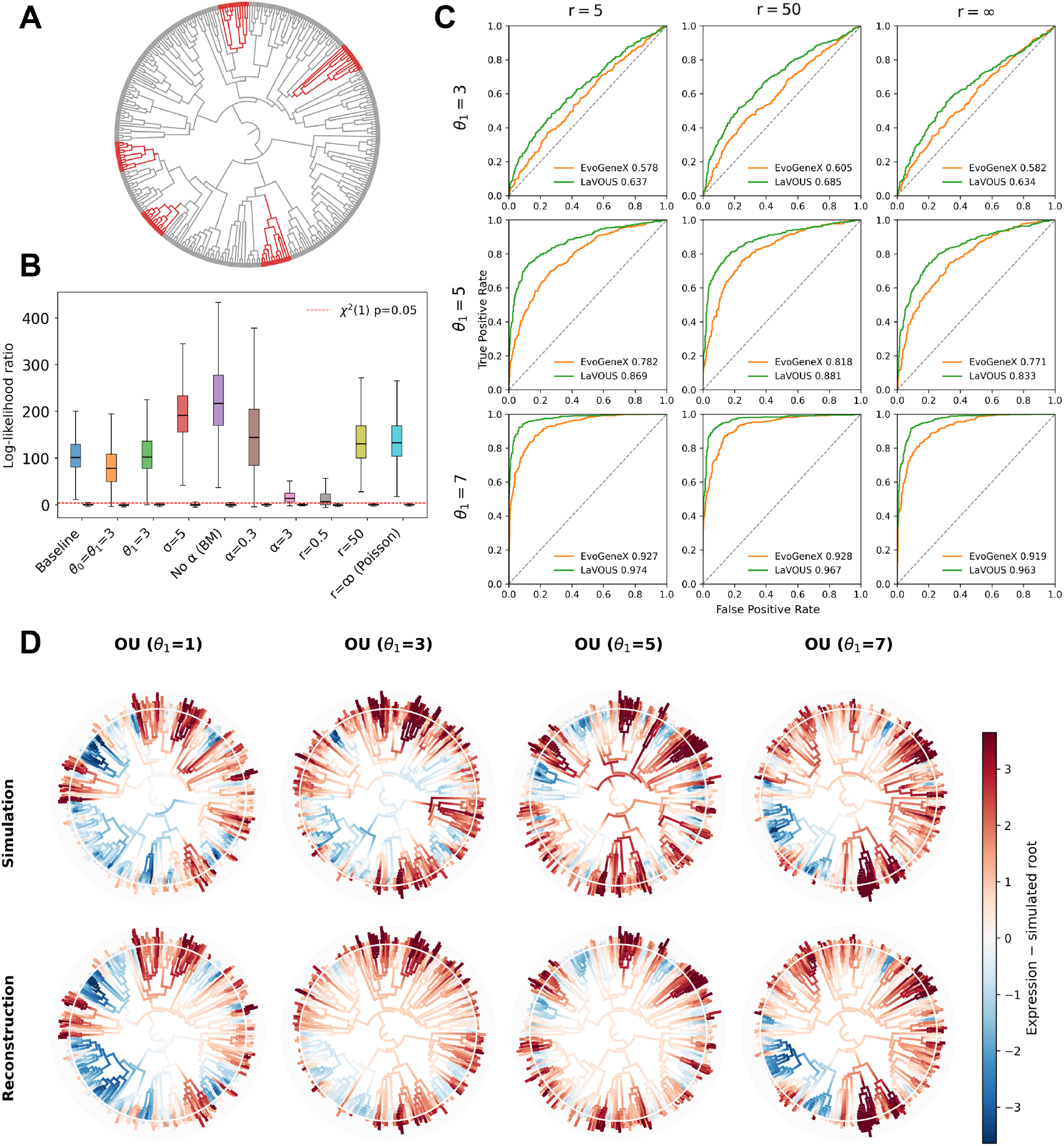
Same simulation experiments as in Fig. 2 using a permutated tree structure. The tree was permutated by randomly swapping branches in Fig. 2A, resulting in different tree structures and regime labels.

**Figure S2:**
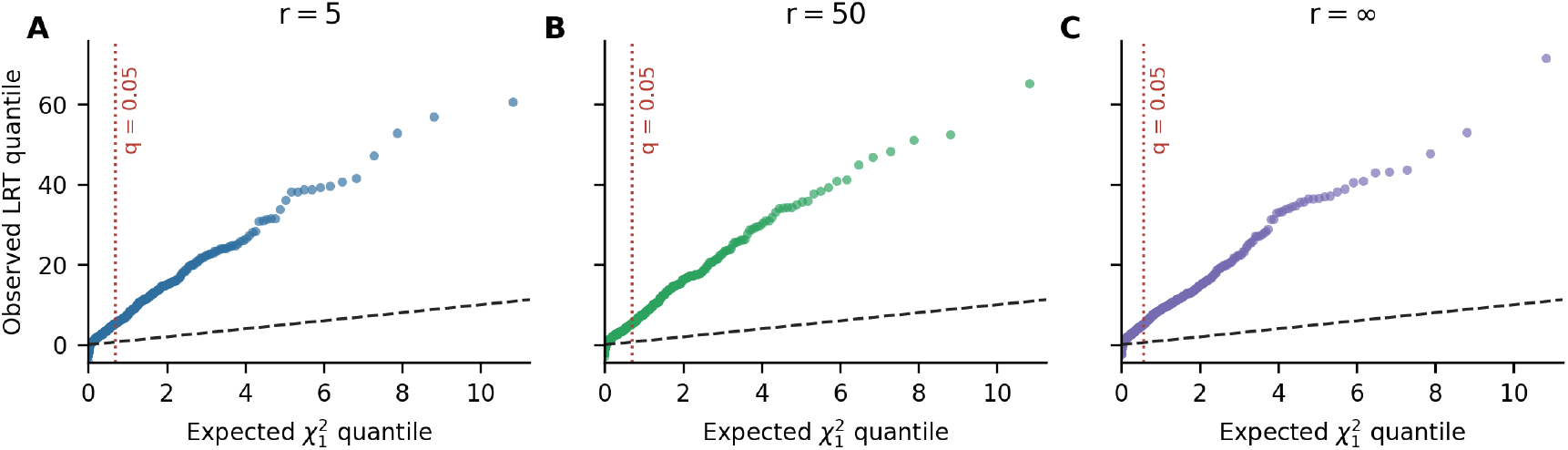
Null likelihood-ratio statistics compared with the asymptotic *χ*^2^ reference. QQ plots show the per-gene LaVOUS differential-test LRTs for the three *θ*_0_ = *θ*_1_ null simulation settings in Fig. 2C. The dashed line in black indicates equality with a *χ*^2^ distribution with one degree of freedom. The vertical dashed line in red indicates the threshold of *q* = 0.05 in *χ*^2^ after Benjamini-Hochberg correction. In all three null settings, the observed LRTs show strong upward tail deviation from the chi-square reference, explaining why *χ*^2^-calibrated *p*-values are anti-conservative for these simulations.

**Figure S3:**
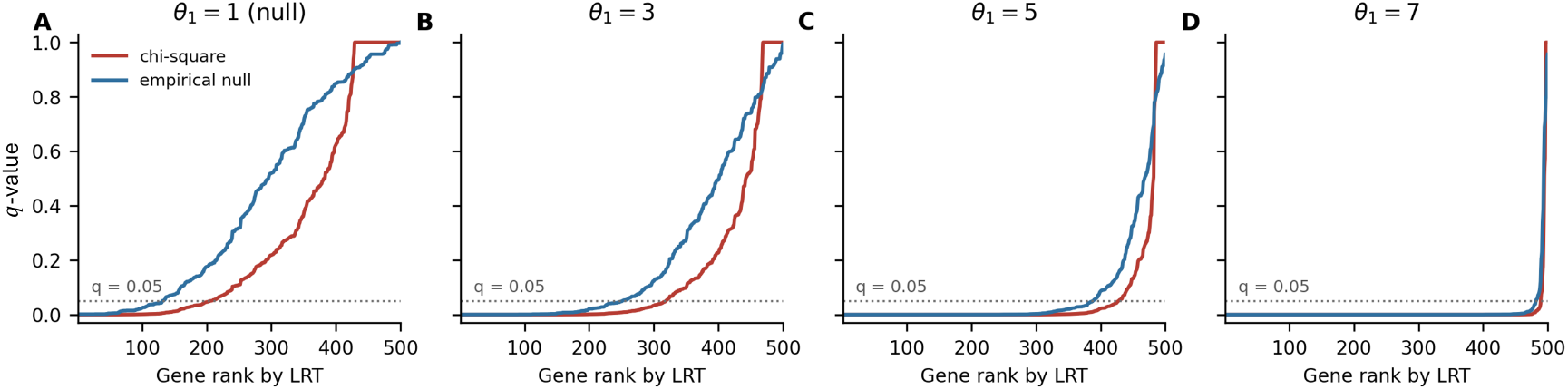
The *q*-value distributions under *χ*^2^ and empirical-null calibration using example *r* = 5 simulations from Fig. 2C. Genes were ordered by decreasing LaVOUS LRT and the dotted horizontal line marks *q* = 0.05. The empirical-null calibration uses the pooled null from the 10,000 simulating samples and shifts *q*-values relative to the asymptotic *χ*^2^ calibration, especially for weaker effects.

**Figure S4:**
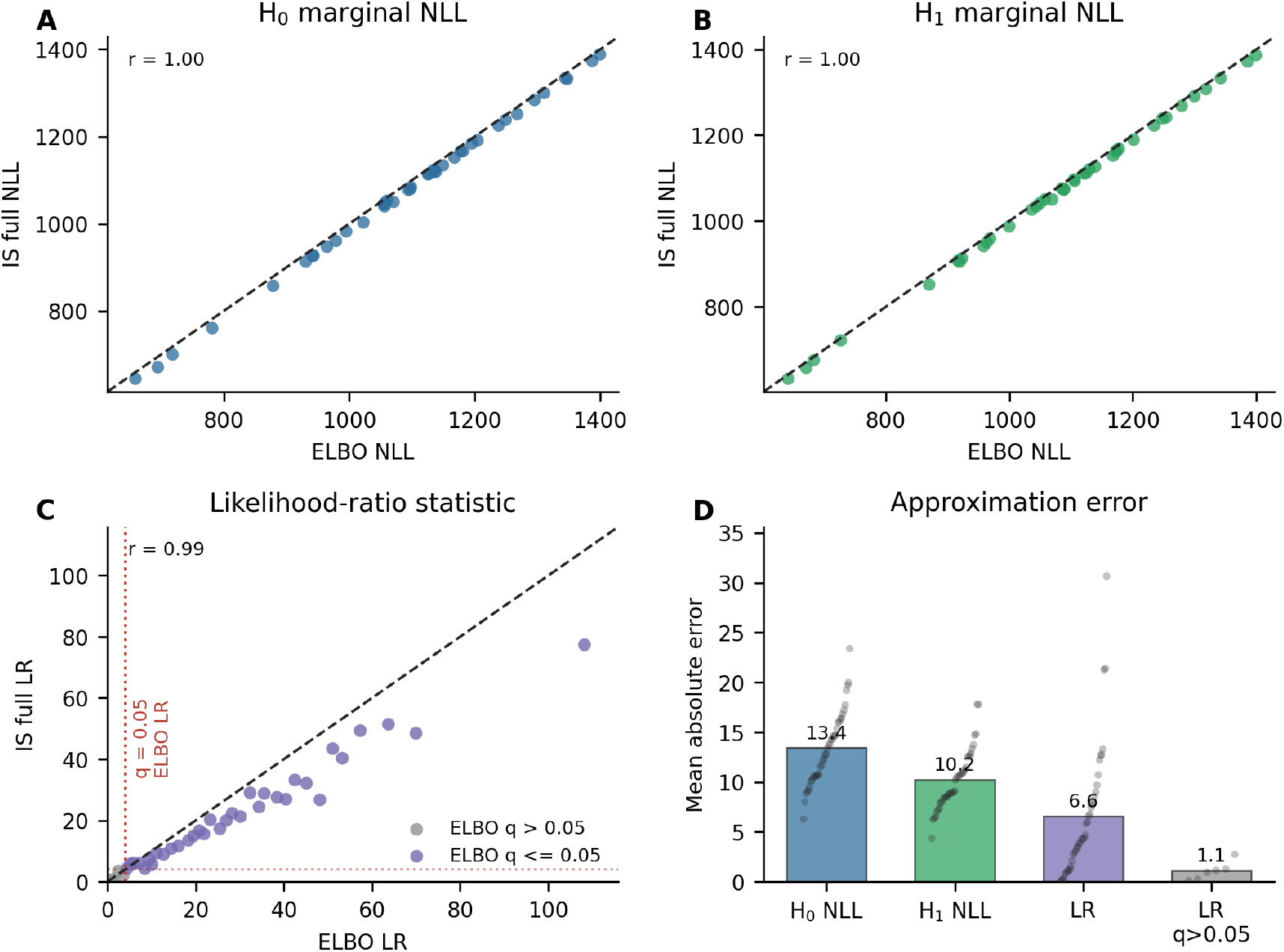
ELBO approximation quality assessed by importance-sampling (IS) marginal likelihood estimates. IS used an overdispersed-q proposal, 0.8 *q*(*z*|*x*) + 0.2 *q*_2_(*z*|*x*), where *q*_2_ has the same mean as q and 2 times the posterior standard deviation, with 3 independent replicates of *S* = 512 samples per gene and hypothesis (1536 pooled samples total). Panels A and B compare the saved variational negative ELBOs with IS-estimated full marginal negative log likelihoods (NLLs) for *H*_0_ and *H*_1_, respectively, using a stratified subset of 40 genes from the *θ*_1_ = 5, *r* = 5 simulation from Fig. 2C. Because the original differential-test run excluded constant terms, the gene-wise negative-binomial observation constant was added back to the saved ELBO losses before these absolute NLL comparisons. Panel C compares the ELBO-based likelihood-ratio (LR) statistics with the corresponding IS-estimated full LRs. Panel D compares mean absolute error (MAE) for the *H*_0_ NLL, *H*_1_ NLL, and LR, showing that the ELBO approximation is closer to the IS estimate after the subtraction used in the LRT. Moreover, inflation errors mostly occur at very large LRs but have little effect on gene significance.

**Figure S5:**
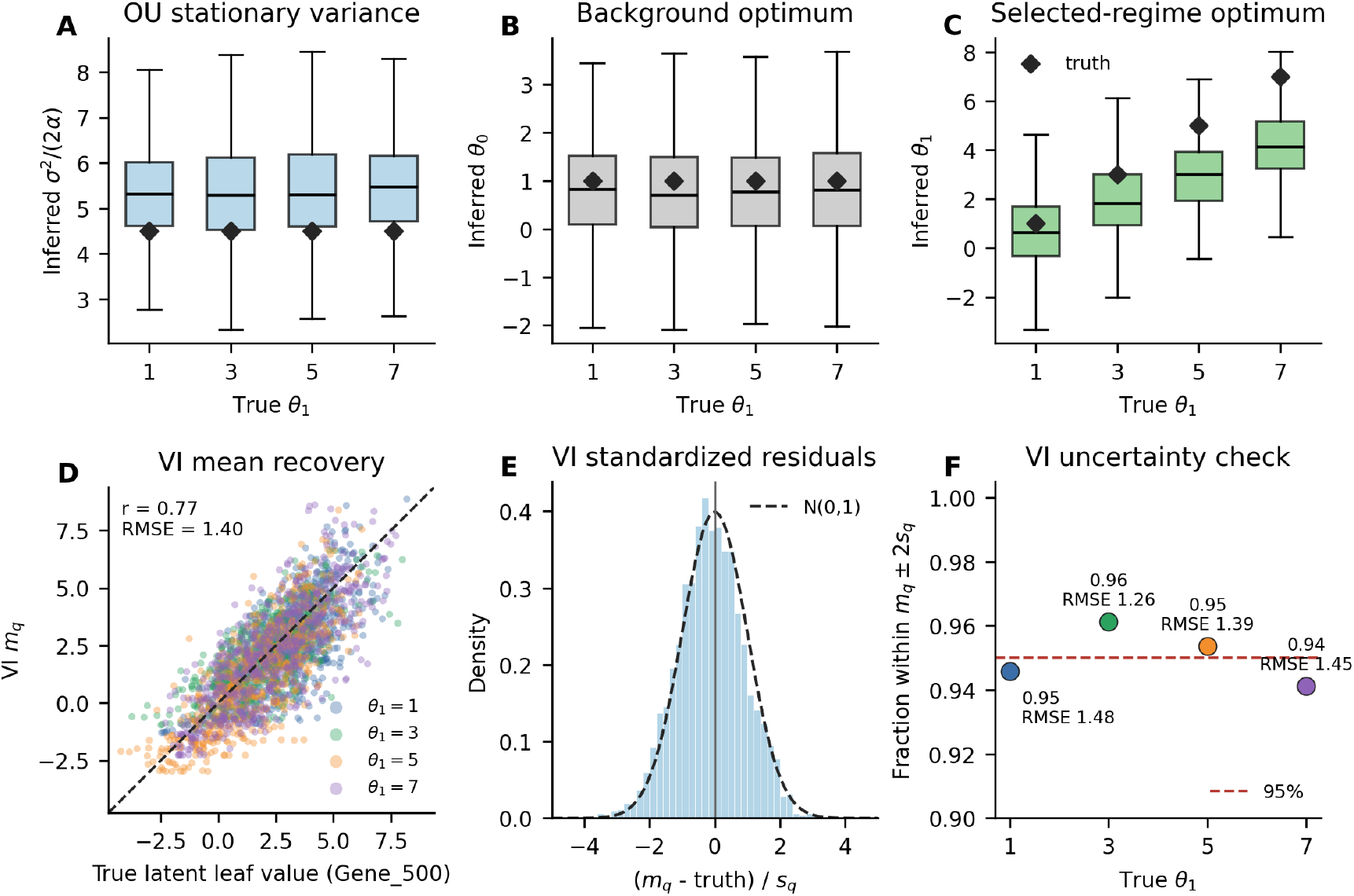
Recovery of OU and variational inference (VI) parameters in simulated data with *r* = 5 from Fig. 2C. Panels A–C summarize *H*_1_ OU parameter estimates across 500 genes from the simulations. Black diamonds mark the simulation truth. Panels D–F evaluate VI leaf-level parameters for Gene 500 for which the simulated expression history was stored. Because the simulator writes terminal expression after the softplus transform, true terminal latent values were recovered with the inverse softplus before comparison with VI m_q_. Panel E standardizes residuals by VI *s*_*q*_, and Panel F shows the fraction of leaves covered by *m*_*q*_ *±* 2*s*_*q*_.

**Figure S6:**
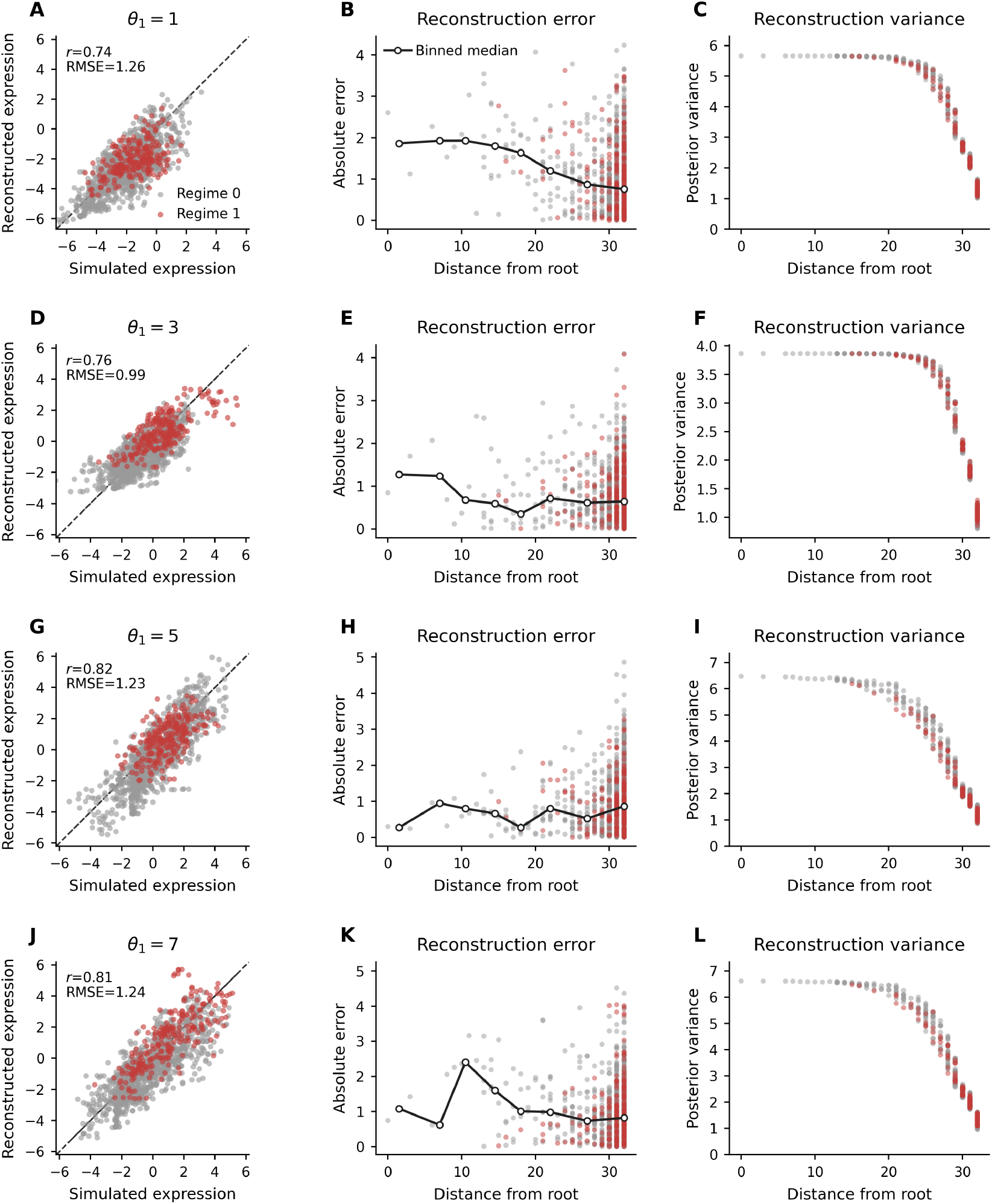
Comparison of reconstructed and simulated expression history. Panel A compares LaVOUS posterior mean reconstructed expression with the simulated latent expression at matched tree nodes for the Gene 500 OU example with *θ*_1_ = 1, *r* = 5 from Fig. 2D. Values are centered on the simulated root for each scenario, matching the color centering used in Fig. 2D. Gray points are nodes from background branches and red points are nodes from selected branches; dashed lines indicate equality between simulated and reconstructed expression. Panel B shows reconstruction error along tree branches from the root for the same example. Panel C shows reconstruction variance of each node along the tree from the root. D–F, same analysis for *θ*_1_ = 3, *r* = 5. G–I, same analysis for *θ*_1_ = 5, *r* = 5. J–L, same analysis for *θ*_1_ = 7, *r* = 5.

## Notes

https://github.com/Jiawei-Xing/LaVOUS

### Competing Interest Statement

The authors have declared no competing interest.

### Summary of Updates

Supplementary figures updated; minor changes to the main text.

## References

[1] Salvador-Martínez, I., Grillo, M., Averof, M. & Telford, M. J. Is it possible to reconstruct an accurate cell lineage using CRISPR recorders? Elife 8, e40292 (2019).

[2] McKenna, A. et al. Whole-organism lineage tracing by combinatorial and cumulative genome editing. Science 353, aaf7907 (2016).

[3] Kalhor, R. et al. Developmental barcoding of whole mouse via homing CRISPR. Science 361, eaat9804 (2018).

[4] Wagner, D. E. & Klein, A. M. Lineage tracing meets single-cell omics: opportunities and challenges. Nature Reviews Genetics 21, 410–427 (2020).

[5] VanHorn, S. & Morris, S. A. Next-generation lineage tracing and fate mapping to interrogate development. Developmental Cell 56, 7–21 (2021).

[6] Jones, M. G., Yang, D. & Weissman, J. S. New tools for lineage tracing in cancer in vivo. Annual Review of Cancer Biology 7, 111–129 (2023).

[7] Zhang, X., Huang, Y., Yang, Y.Wang, Q.-E. & Li, L. Advancements in prospective single-cell lineage barcoding and their applications in research. Genome Research 34, 2147 (2024).

[8] Alemany, A., Florescu, M., Baron, C. S., Peterson-Maduro, J. & Van Oudenaarden, A. Whole-organism clone tracing using single-cell sequencing. Nature 556, 108–112 (2018).

[9] Spanjaard, B. et al. Simultaneous lineage tracing and cell-type identification using CRISPR–Cas9-induced genetic scars. Nature Biotechnology 36, 469–473 (2018).

[10] Raj, B. et al. Simultaneous single-cell profiling of lineages and cell types in the vertebrate brain. Nature Biotechnology 36, 442–450 (2018).

[11] Trapnell, C. et al. The dynamics and regulators of cell fate decisions are revealed by pseudotemporal ordering of single cells. Nature Biotechnology 32, 381–386 (2014).

[12] Haghverdi, L., Büttner, M., Wolf, F. A., Buettner, F. & Theis, F. J. Diffusion pseudotime robustly reconstructs lineage branching. Nature Methods 13, 845–848 (2016).

[13] Street, K. et al. Slingshot: cell lineage and pseudotime inference for single-cell transcriptomics. BMC Genomics 19, 477 (2018).

[14] Chan, M. M. et al. Molecular recording of mammalian embryogenesis. Nature 570, 77–82 (2019).

[15] Quinn, J. J. et al. Single-cell lineages reveal the rates, routes, and drivers of metastasis in cancer xenografts. Science 371, eabc1944 (2021).

[16] Simeonov, K. P. et al. Single-cell lineage tracing of metastatic cancer reveals selection of hybrid EMT states. Cancer Cell 39, 1150–1162 (2021).

[17] Yang, D. et al. Lineage tracing reveals the phylodynamics, plasticity, and paths of tumor evolution. Cell 185, 1905–1923.e25 (2022).

[18] Zhu, H., Liu, X., Tang, M., Lui, K. O. & Zhou, B. Lineage tracing from cellular heritage to disease destiny. Nature Genetics 1–15 (2026).

[19] Jones, M. G. et al. Inference of single-cell phylogenies from lineage tracing data using Cassiopeia. Genome Biology 21, 92 (2020).

[20] Chu, G., Mai, U., Schmidt, H. & Raphael, B. J. Maximum likelihood inference of time-scaled cell lineage trees with mixed-type missing data using LAML. Genome Biology 26, 189 (2025).

[21] Staklinski, S. J. et al. Bayesian inference of tissue-migration histories in metastatic cancer from cell-lineage tracing data. Cell Genomics (2026).

[22] Siepel, A., Hassett, R. & Staklinski, S. J. VINE: Variational inference for scalable Bayesian reconstruction of species and cell-lineage phylogenies. bioRxiv (2025).

[23] Wang, K., He, X. & Hu, Z. Computational approaches for multimodal lineage tracing. Nature Reviews Genetics 1–18 (2026).

[24] Kang, Z., Chen, H., Li, S.Wang, S.-W. & Zhou, B. Evolving strategies for lineage tracing: Genetic markers, synthetic barcodes, and natural variants. Cell Stem Cell (2026).

[25] Forrow, A. & Schiebinger, G. LineageOT is a unified framework for lineage tracing and trajectory inference. Nature Communications 12, 4940 (2021).

[26] Wang, K. et al. PhyloVelo enhances transcriptomic velocity field mapping using monotonically expressed genes. Nature Biotechnology 42, 778–789 (2024).

[27] Wang, S.-W., Herriges, M. J., Hurley, K., Kotton, D. N. & Klein, A. M. CoSpar identifies early cell fate biases from single-cell transcriptomic and lineage information. Nature Biotechnology 40, 1066–1074 (2022).

[28] Ouardini, K. et al. Reconstructing unobserved cellular states from paired single-cell lineage tracing and transcriptomics data. bioRxiv (2021).

[29] Majima, K. et al. LineageVAE: reconstructing historical cell states and transcriptomes toward unobserved progenitors. Bioinformatics 40, btae520 (2024).

[30] Schiffman, J. S. et al. Defining heritability, plasticity, and transition dynamics of cellular phenotypes in somatic evolution. Nature Genetics 56, 2174–2184 (2024).

[31] DeTomaso, D. & Yosef, N. Hotspot identifies informative gene modules across modalities of single-cell genomics. Cell Systems 12, 446–456 (2021).

[32] Schlüter, H. M. & Uhler, C. Integrating representation learning, permutation, and optimization to detect lineage-related gene expression patterns. Nature Communications 16, 1062 (2025).

[33] Felsenstein, J. Phylogenies and the comparative method. The American Naturalist 125, 1–15 (1985).

[34] Hansen, T. F. Stabilizing selection and the comparative analysis of adaptation. Evolution 51, 1341–1351 (1997).

[35] Butler, M. A. & King, A. A. Phylogenetic comparative analysis: a modeling approach for adaptive evolution. The American Naturalist 164, 683–695 (2004).

[36] Bedford, T. & Hartl, D. L. Optimization of gene expression by natural selection. Proceedings of the National Academy of Sciences 106, 1133–1138 (2009).

[37] Brawand, D. et al. The evolution of gene expression levels in mammalian organs. Nature 478, 343–348 (2011).

[38] Rohlfs, R. V., Harrigan, P. & Nielsen, R. Modeling gene expression evolution with an extended Ornstein–Uhlenbeck process accounting for within-species variation. Molecular Biology and Evolution 31, 201–211 (2014).

[39] Chen, J. et al. A quantitative framework for characterizing the evolutionary history of mammalian gene expression. Genome Research 29, 53–63 (2019).

[40] Pal, S., Oliver, B. & Przytycka, T. M. Stochastic modeling of gene expression evolution uncovers tissue- and sex-specific properties of expression evolution in the drosophila genus. Journal of Computational Biology 30, 21–40 (2023).

[41] Hirsch, M. G. et al. Stochastic modeling of single-cell gene expression adaptation reveals non-genomic contribution to evolution of tumor subclones. Cell Systems 16, 101156 (2025).

[42] Stuart, H. & McKenna, A. SCOUT: Ornstein–Uhlenbeck modelling of gene expression evolution on single-cell lineage trees. bioRxiv (2025).

[43] Sarkar, A. & Stephens, M. Separating measurement and expression models clarifies confusion in single-cell RNA sequencing analysis. Nature Genetics 53, 770–777 (2021).

[44] Lopez, R., Regier, J., Cole, M. B., Jordan, M. I. & Yosef, N. Deep generative modeling for single-cell transcriptomics. Nature methods 15, 1053–1058 (2018).

[45] Pagel, M. Inferring evolutionary processes from phylogenies. Zoologica Scripta 26, 331–348 (1997).

[46] Pagel, M. Inferring the historical patterns of biological evolution. Nature 401, 877–884 (1999).

[47] Freckleton, R. P., Harvey, P. H. & Pagel, M. Phylogenetic analysis and comparative data: a test and review of evidence. The American Naturalist 160, 712–726 (2002).

[48] El-Kebir, M., Satas, G. & Raphael, B. J. Inferring parsimonious migration histories for metastatic cancers. Nature Genetics 50, 718–726 (2018).

[49] Matouk, I. J. et al. The role of the oncofetal H19 lncRNA in tumor metastasis: orchestrating the EMT-MET decision. Oncotarget 7, 3748 (2015).

[50] Deuel, T. F., Zhang, N., Yeh, H.-J., Silos-Santiago, I. & Wang, Z.-Y. Pleiotrophin: a cytokine with diverse functions and a novel signaling pathway. Archives of Biochemistry and Biophysics 397, 162–171 (2002).

[51] Lindström, M. S. NPM1/B23: a multifunctional chaperone in ribosome biogenesis and chromatin remodeling. Biochemistry Research International 2011, 195209 (2011).

[52] Fyfe-Desmarais, G., Desmarais, F., Rassart, É . & Mounier, C. Apolipoprotein D in oxidative stress and inflammation. Antioxidants 12, 1027 (2023).

[53] Arensdorf, A. M. & Thomas Rutkowski, D. Endoplasmic reticulum stress impairs IL-4/IL-13 signaling through C/EBPβ-mediated transcriptional suppression. Journal of Cell Science 126, 4026–4036 (2013).

[54] Chang, C., Worley, B. L., Phaëton, R. & Hempel, N. Extracellular glutathione peroxidase GPx3 and its role in cancer. Cancers 12, 2197 (2020).

[55] Jiang, X., Zhang, J. & Huang, Y. Tetraspanins in cell migration. Cell Adhesion & Migration 9, 406–415 (2015).

[56] de Assis, F. L. et al. Tracking B cell responses to the SARS-CoV-2 mRNA-1273 vaccine. Cell Reports 42 (2023).

[57] Weber, L. L. et al. Isotype-aware inference of B cell clonal lineage trees from single-cell sequencing data. Cell Genomics 4 (2024).

[58] Hoehn, K. B., Pybus, O. G. & Kleinstein, S. H. Phylogenetic analysis of migration, differentiation, and class switching in B cells. PLoS Computational Biology 18, e1009885 (2022).

[59] Krammer, F. The human antibody response to influenza A virus infection and vaccination. Nature Reviews Immunology 19, 383–397 (2019).

[60] Stavnezer, J. & Schrader, C. E. IgH chain class switch recombination: mechanism and regulation. The Journal of Immunology 193, 5370–5378 (2014).

[61] Horns, F. et al. Lineage tracing of human B cells reveals the in vivo landscape of human antibody class switching. Elife 5, e16578 (2016).

[62] Reimold, A. M. et al. Plasma cell differentiation requires the transcription factor XBP-1. Nature 412, 300–307 (2001).

[63] Kumar, N., Arthur, C. P., Ciferri, C. & Matsumoto, M. L. Structure of the secretory immunoglobulin A core. Science 367, 1008–1014 (2020).

[64] Xie, L. et al. Comprehensive spatiotemporal mapping of single-cell lineages in developing mouse brain by CRISPR-based barcoding. Nature Methods 20, 1244–1255 (2023).

[65] Bertoldi, M. Mammalian Dopa decarboxylase: structure, catalytic activity and inhibition. Archives of Biochemistry and Biophysics 546, 1–7 (2014).

[66] Wolf, F. A., Angerer, P. & Theis, F. J. SCANPY: large-scale single-cell gene expression data analysis. Genome Biology 19, 15 (2018).

[67] Andersson, E. R. et al. Wnt5a regulates ventral midbrain morphogenesis and the development of A9–A10 dopaminergic cells in vivo. PloS One 3, e3517 (2008).

[68] Yagi, T. & Takeichi, M. Cadherin superfamily genes: functions, genomic organization, and neurologic diversity. Genes & Development 14, 1169–1180 (2000).

[69] McInnes, L., Healy, J. & Melville, J. UMAP: Uniform Manifold Approximation and Projection for Dimension Reduction. arXiv preprint arXiv:1802.03426 (2018).

[70] Kolberg, L. et al. g:Profiler—interoperable web service for functional enrichment analysis and gene identifier mapping (2023 update). Nucleic Acids Research 51, W207–W212 (2023).

